# Single-cell transcriptomics yield insights into the stress-immune interplay and inform disease risk

**DOI:** 10.1101/2025.06.05.658150

**Authors:** Sofia Benavides, Oksana Kosyk, Lillian Carpenter, Christina Georgiou, Brian C. Miller, Natalie Stanley, Anthony S. Zannas

## Abstract

Environmental stress contributes to several disease conditions – including cardiovascular disease, mental illness, and autoimmune disorders – through immune dysregulation. Stress operates through the hypothalamic-pituitary-adrenal and sympathoadrenal axes, leading to peripheral secretion of cortisol and norepinephrine, respectively. While stress profoundly influences immune function, the underlying single-cell and cell-type-specific mechanisms remain poorly understood. To address this knowledge gap, we employed an ex vivo model investigating the transcriptomic responses of primary peripheral blood mononuclear cells and neutrophils to physiological stress levels (i.e., levels reached during in vivo stress) of cortisol and norepinephrine at single-cell resolution. We identified novel stress-hormone-dependent and cell-type-specific transcriptomic changes, including uniquely regulated genes and inflammation-related signatures within distinct innate (e.g., *CD163*^+^ monocytes/macrophages, neutrophils) and other immune cell types. The single-cell-derived signatures, applied to two independent human cohorts, effectively distinguished stress-related diseases. Our findings provide cell-type-level insights into how stress hormones modulate immune function relevant to stress-related conditions.

## INTRODUCTION

Environmental stress, an unavoidable characteristic of every society (*1*), has been linked to an increased risk for various diseases (*2–19*). For example, studies have established an association between excessive psychosocial stress and an increased risk of cardiovascular disease (*2–7*), mental illness (*20, 21*), and autoimmune disorders (*13–16*). While the underlying mechanisms are poorly understood, immune dysregulation has been repeatedly highlighted as a key process shared across stress-related conditions (*22–28*).

Stress operates through two distinct biological pathways: the Hypothalamic-Pituitary-Adrenal (HPA) axis (*29*) and the sympathoadrenal axis (*30*). These pathways culminate in the secretion of stress hormones – cortisol and norepinephrine – into the periphery, where they interact with immune cells (*31*). Controversy exists regarding the effects of stress hormones on immune cells (*32*); some literature suggests an anti-inflammatory response (*33, 34*), while other studies demonstrate pro-inflammatory effects (*22, 35*). However, to our knowledge, all previous research on stress hormones and immune cells has focused on analyzing a mixture of cells (i.e., bulk-level analysis)(*36–39*), thereby not accounting for cell type-specific effects that could play a critical role in mediating stress pathology. Additionally, most studies investigating stress-induced immune dysfunction have used in-vivo models. While these models are critical for studying complex effects on body systems, they are confounded by natural shifts in cell populations and changes in cell-type distribution that constantly occur in vivo, thus providing limited single-cell and cell-type-specific insights into how stress regulates immune function.

To address these knowledge gaps, here we employ a cell model (ex vivo) approach and single-cell RNA sequencing (scRNA-seq) to directly assess the immune effects of physiological stress hormone exposure at the single-cell level. To model physiological stress in culture, primary peripheral blood mononuclear cells (PBMCs) and neutrophils isolated from human donors were exposed to the endogenous human stress hormones cortisol and norepinephrine at concentrations known to be reached in human tissues during in vivo stress (*40–47*). This approach enabled us to model stress ex vivo and conduct a comprehensive analysis of the unique immune transcriptomic signatures of physiological stress hormone exposure at the single-cell and cell-type levels. The single-cell-derived signatures, applied to two independent human cohorts, effectively distinguished stress-related diseases. Our findings provide new insights into how stress hormones influence immune cell function and the immune cell-type-specific mechanisms underlying stress-related conditions.

## RESULTS

### Immune single-cell transcriptional profiling after ex vivo exposure to physiological stress hormone levels

To evaluate the effects of stress hormones on immune cell function, we obtained whole blood samples from four male and four female healthy donors and isolated PBMCs and neutrophils. Cells from each donor were treated with vehicle control (0.0001% DMSO), 100 nM cortisol (CORT), or 10 nM norepinephrine (NE) (Fig. 1A). These concentrations were selected based on previous literature indicating that glucocorticoids can reach tissue concentrations of 100 nM (*40–44*), and norepinephrine reaches 10 nM circulating concentration (*45–47*) during in vivo stress exposure. This robust experimental design enables within-donor comparisons, thereby accounting for potential inter-individual variability in transcriptional responses.

**Fig. 1.**
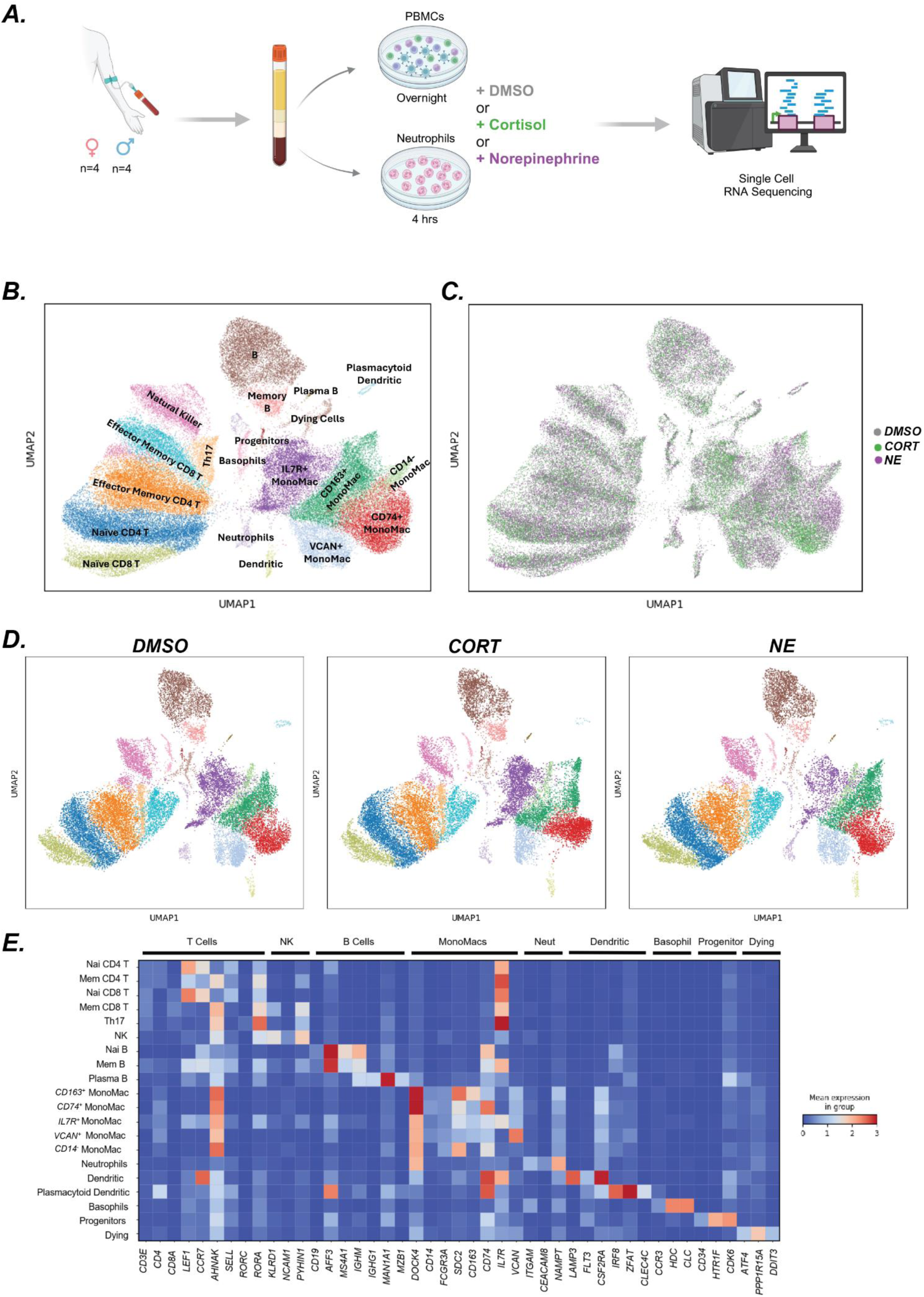
Single-cell RNA sequencing analysis of immune cells after ex vivo exposure to stress hormones. (A) Graphical representation of experimental model. PBMCs and neutrophils from healthy human donors were isolated and exposed to control (DMSO) and treatment (cortisol (CORT), norepinephrine (NE)) groups. After 4 hours of incubation for neutrophils (due to their shorter lifespan (*85*)) and overnight incubation for PBMCs, cells were prepared for single-cell RNA sequencing. (B) UMAP plot showing 62,975 cells from all conditions, colored by cell type. (C) UMAP plot of all cells colored by treatment group (DMSO, CORT, NE). (D) UMAP plots separated by treatment group, highlighting that all cell type clusters are represented in each condition. (E) Heatmap displaying differentially expressed genes (DEGs), canonical marker genes, and genes identified in previous PBMC studies used for cell type annotation.

After excluding low-quality cells and potential doublets, we retained 62,975 high-quality cells: 23,000 from the vehicle, 20,707 from the CORT, and 19,268 from the NE group. Quality control metrics for the retained cells are shown in Fig. S1B–F. Cluster analysis identified 20 distinct cell clusters, visualized using a standard uniform manifold approximation and projection (UMAP) approach (*48*) (Fig. 1B). Although the treatment groups showed similar representation across all 20 clusters, initial UMAP visualization by treatment revealed subtle shifts in cell distribution attributable to the treatments (Figs. 1C, D).

The clusters encompassed all major cell types typically found in peripheral blood, including lymphocytes, monocytes, and neutrophils. The distribution of cells by individual sample is shown in Fig. S1A. All clusters were successfully annotated using the top differentially expressed genes (DEGs), canonical markers, and reference data from previously published scRNA-seq datasets (*49, 50*) (Fig. 1E).

### Exposure to physiological stress hormone levels alters immune cell type composition

While stress-induced changes in immune cell type composition have been well-documented in vivo, the functional impact of CORT and NE on distinct immune cell types remains unclear. Addressing this question in culture allows for evaluating the direct effects of stress hormones on immune cell composition, free from the confounding effects of cell mobilization and redistribution observed in vivo. Using density plots, we qualitatively observed treatment-dependent changes in immune composition. CORT treatment resulted in a general decrease in lymphocyte proportions and altered the distribution of monocyte/macrophage (MonoMac) populations (Fig. 2A). In contrast, NE treatment produced a distinct pattern of T cell distributions and a unique MonoMac distribution that differed from the control and CORT-treated groups (Fig. 2A).

**Fig. 2.**
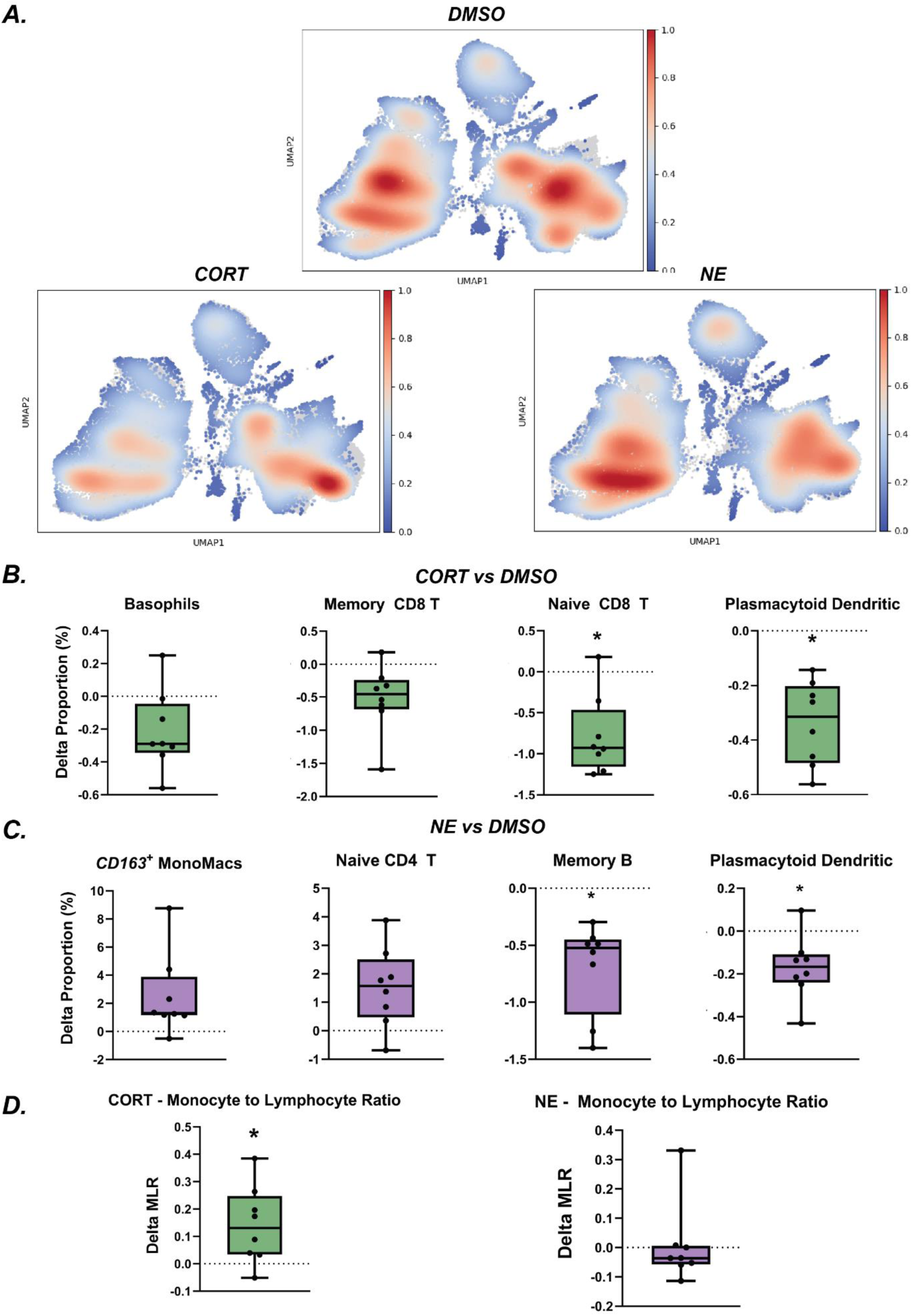
Stress hormone treatment induces treatment-specific immune remodeling. (A) Density plots showing cell type distributions in control and treatment groups. Blue represents lower cell density, while red indicates higher density. (B) Changes in the percentages of plasmacytoid dendritic cells, naïve CD8 T cells, memory CD8 T cells, and basophils following CORT exposure. While all cell types shown were nominally significant, only naïve CD8 T cells and plasmacytoid dendritic cells remained significant after multiple testing correction using the Bonferroni method. Graphs depict the deltas between CORT and DMSO. Paired t-tests were used to test for significance (*p < 0.0025). (C) Changes in the percentages of plasmacytoid dendritic cells, naïve CD4 T cells, memory B cells, and CD163^+^ MonoMacs following NE exposure. While all cell types shown were nominally significant, only memory B cells and plasmacytoid dendritic cells remained significant after multiple testing correction using the Bonferroni method. Graphs show the deltas between NE and DMSO. Paired t-tests were used to test for significance (*p < 0.0025). (D) Differences in the MLR between treatment and control groups. Graphs represent the deltas for CORT (green) and NE (purple) compared to control. Paired t-tests were used to test for significance (*p < 0.05).

We further quantified these changes, leveraging the strength of our experimental design that enables within-donor comparisons between control and treated cells. These comparisons revealed significant stress hormone-induced changes in cell type proportions. Specifically, CORT treatment led to nominally significant reductions in memory and naïve CD8 T cells, basophils, and plasmacytoid dendritic cells. However, after correcting for multiple testing, only the decreases in naïve CD8 T cells and plasmacytoid dendritic cells remained statistically significant (Fig. 2B). Similarly, NE treatment resulted in nominally significant decreases in plasmacytoid dendritic cells and memory B cells, as well as increases in naïve CD4 T cells and *CD163*+ MonoMac populations (Fig. 2C). After multiple testing corrections, only the reductions in memory B cells and plasmacytoid dendritic cells remained statistically significant. Additional patterns of change in cell type proportions were observed, but did not reach statistical significance (fig. S2A-B).

To validate these findings on cell type composition, we performed RNAScope analysis in four independent donors (two males and two females). We chose to focus on validating the CORT-driven decrease in naïve CD8⁺ T cells due to the greater abundance of this cell type among the statistically significant cell populations (naïve CD8 T ∼6.26%, memory B ∼1.5%, plasmacytoid dendritic cells ∼0.29%), which allowed for greater statistical power. We observed a pattern of decreased naïve CD8⁺ T cells across donors (fig. S3A-C), but the overall result did not reach statistical significance. This may reflect limitations in power and the lower sensitivity of the RNAScope assay compared to scRNA-seq.

To further evaluate the potential clinical relevance of stress hormone-induced immune remodeling, we calculated the monocyte-to-lymphocyte ratio (MLR), an established prognostic marker of conditions such as kidney disease (*51*), cardiovascular mortality (*52*), and cancer (*53, 54*). Our analysis revealed a significant increase in MLR following CORT, but not NE, treatment (Fig. 2D). These findings indicate that, even in the absence of cell mobilization and redistribution, exposure to physiological stress hormone levels can selectively remodel the immune cell composition.

### Exposure to physiological stress hormone levels induces cell type-specific gene expression changes

Next, we investigated the extent to which exposure to CORT and NE concentrations corresponding to physiological levels of stress alters immune cell-type-specific gene expression.

CORT increased the expression of well-characterized glucocorticoid-responsive genes, such as *FKBP5* and *ZBTB16* (Fig. 3A), at the bulk level (i.e., when pooling all cell types), confirming the effectiveness of our ex vivo treatment. In total, we identified 450 bulk-level differentially expressed genes (DEGs) with an absolute log fold change (LFC) > 0.25 and a false discovery rate (FDR)-corrected *p*-value (*q*) < 0.05. Of these, 251 genes were downregulated and 199 were upregulated (table S1). We validated several top DEGs (*VSIG4*, *FKBP5*, *ZBTB16*) using qPCR (Fig. 3C). We further examined the expression patterns of the top DEGs due to CORT treatment across different immune cell types. Notably, CORT induced both shared and distinct transcriptional changes across cell types (Fig. 3B). For instance, *VSIG4* was markedly upregulated in progenitor cells (LFC = 24.4), while it was strongly downregulated in plasmacytoid dendritic cells (LFC = −23.4). Among all DEGs, *FKBP5* was the only gene consistently upregulated across all cell types due to CORT treatment (with an absolute LFC > 0.25). Although not part of the top 10 DEGs, *RNU4-2* showed consistent upregulation across most cell types except plasmacytoid dendritic cells, while *CCND3* exhibited consistent upregulation across all cell types except plasmacytoid dendritic and progenitor cells.

**Fig. 3.**
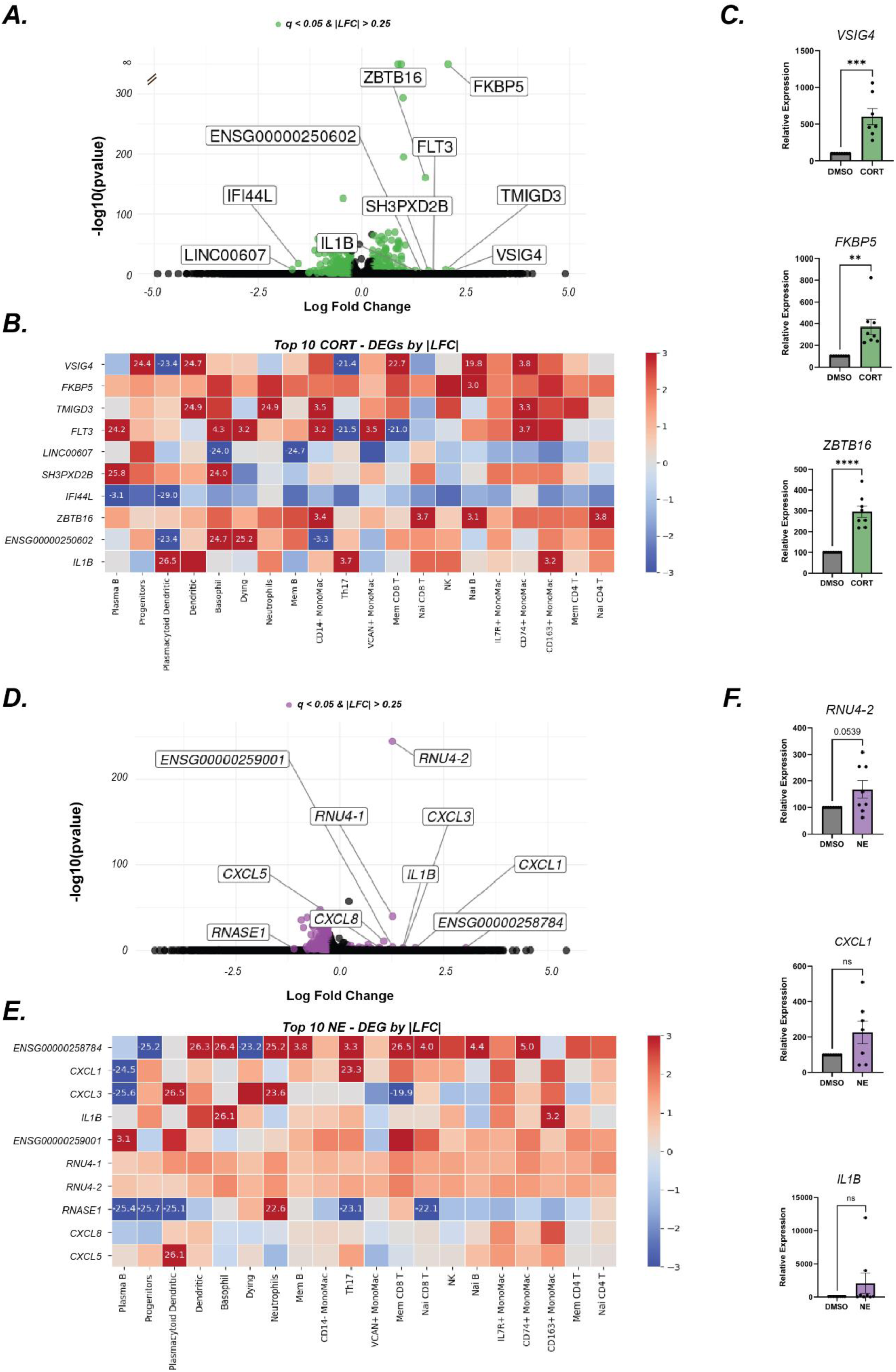
CORT and NE treatment lead to cell-type-specific gene expression changes. (A) Volcano plot showing bulk gene expression changes due to CORT treatment. Green dots represent significant genes with an absolute log fold change (LFC) greater than 0.25 and an FDR-corrected p-value (q) of less than 0.05. The top 10 genes, based on LFC, are labeled. (B) Heatmap of the top 10 differentially expressed genes (DEGs) based on LFC, categorized by cell type. (C) qPCR validation of gene expression changes due to CORT treatment. Statistical significance was determined by Student’s t-test. *p < 0.05, **p < 0.01, ***p < 0.001, ****p < 0.0001. (D) Volcano plot showing bulk gene expression changes due to NE treatment. Purple dots indicate significant genes with an absolute LFC greater than 0.25 and q less than 0.05. The top 10 genes according to LFC are labeled. (E) Heatmap of the top 10 DEGs based on LFC, categorized by cell type (F) qPCR validation of gene expression changes due to NE treatment. Statistical significance was determined by Student’s t-test.

NE exposure led to a total of 472 DEGs with an absolute LFC greater than 0.25 at the bulk level. Strikingly, 451 of these genes were downregulated, while only 21 were upregulated (Fig. 3D, table S2). We validated several top DEGs (*RUN4-2*, *CXCL1, IL1B*) via qPCR, and although these results did not reach statistical significance, they followed a similar expression pattern as our scRNA-seq findings (Fig. 3F). We then analyzed the cell type-specific effects and found that, similar to CORT, NE treatment also induced cell type-specific gene expression patterns. For example, *CXCL3* was strongly upregulated in neutrophils (LFC = 23.59) and plasmacytoid dendritic cells (LFC = 26.51), whereas it was markedly downregulated in Memory CD8 T cells (LFC = −19.92) and Plasma B cells (LFC = −25.61) (Fig. 3E). Among all the DEGs, we identified *RNU4-2* as the single gene consistently upregulated in the same direction across all immune cell types due to NE treatment (with an absolute LFC > 0.25). Given the lack of a benchmark gene marker for assessing NE signaling, our analyses for the first time identify *RNU4-2* as a strong candidate for further investigation in immune cells.

Interestingly, CORT and NE treatments shared 100 common DEGs at the bulk level, of which 62 were differentially regulated in the same direction and 38 were regulated in opposite directions.

To our knowledge, this is the first study showing that stress hormones induce distinct and drastically differential effects on gene expression that are dependent on immune cell type. These results may help explain the inconsistencies observed in the literature regarding the effects of stress hormones on immune function (*22, 32, 33, 55–57*), as prior studies were conducted at the bulk level.

### Innate immune cell types display distinct inflammatory responses when exposed to CORT and NE

Given our interest in the role of stress in disease-related inflammation, we analyzed how CORT and NE treatments influence differential inflammatory gene expression across immune cell types. To achieve this, we utilized AUCell, a tool designed to assess whether a specific input gene set is enriched within the expression profile of each cell (*58*). For our analysis, we utilized the hallmark inflammatory response gene set, consisting of 200 genes originally developed for gene set enrichment analysis (GSEA)(*59*). The distribution of cells by AUCell score is shown in fig. S4A and S4B. Our findings revealed that cells exposed to stress hormones exhibited cell-type-specific differential expression of inflammation-related genes, with MonoMac cell clusters demonstrating the most pronounced changes (Fig. 4A). To quantify these differences, we compared the AUCell scores for each cell type between the control and treatment groups. Following CORT exposure, neutrophils, *IL7R*^+^ MonoMacs, and *CD163*^+^ MonoMacs showed significant enrichment of inflammation-related gene expression, with *CD163*^+^ MonoMacs exhibiting the most pronounced change (Fig.4B). Similarly, NE treatment resulted in inflammation-related gene enrichment in *IL7R*^+^ MonoMacs and *CD163*^+^ MonoMacs, with *CD163*^+^ MonoMacs again showing the greatest change (Fig. 4C). Interestingly, in both treatment conditions, we observed a significant downregulation of inflammation-related genes in *VCAN*^+^ MonoMacs (Fig. 4A and B). This highlights how the effects of different stress hormones can be contrasting even between closely related cell types, such as those in the MonoMac compartment, emphasizing the importance of cell-type-specific analysis.

**Fig. 4.**
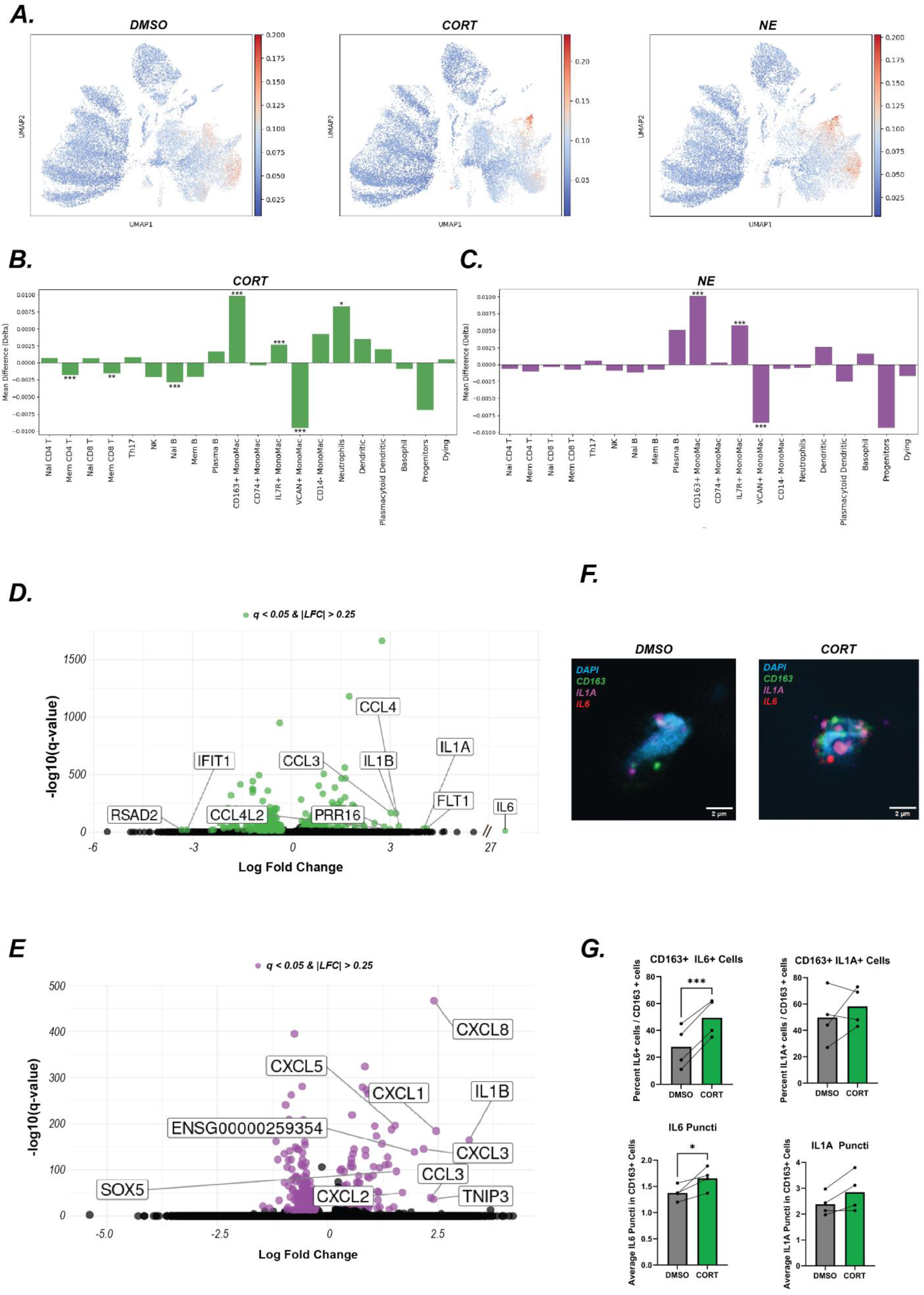
*CD163*^+^ MonoMacs show a distinct inflammatory response to stress hormone treatment. (A) Heatmaps of UMAPs by treatment. Heatmap shows individual cell’s AUCell score depicting inflammatory gene expression enrichment. (B) Quantification of the difference in AUCell enrichment score between CORT and control. Bonferroni– corrected p values: *p < 0.05, **p < 0.01, ***p < 0.001. (C) Quantification of the difference in AUCell enrichment score between NE and control. Bonferroni–corrected p values: ***p < 0.001. (D) Volcano plot showing gene expression changes due to CORT treatment in *CD163*^+^ MonoMacs. Green dots represent significant genes with an absolute log fold change (LFC) greater than 0.25. The top 10 genes, based on LFC, are labeled. (E) Volcano plot showing gene expression changes due to NE treatment in *CD163*^+^ MonoMacs. Green dots represent significant genes with an absolute log fold change (LFC) greater than 0.25. The top 10 genes, based on LFC, are labeled. (F) Representative images of RNAScope analysis. Cells were selected for *CD163*+ expression (green) and then *IL6* (red) and *IL1A* (magenta) puncta were analyzed. (G) Image quantifications of RNAScope analysis. Significance was established using student’s t-test. : *p < 0.05, **p < 0.01, ***p < 0.001.

#### Characterization and validation of CD163+ MonoMacs as a novel stress-activated innate immune cell subtype

Given the observed robust inflammatory signature enrichment in *CD163*^+^ MonoMacs (F-statistic: 75.45, p-value: 4.02e-33) (fig. S4C), we further characterized the stress-driven DEGs specifically in this cell type. CORT exposure resulted in a total of 477 DEGs with an absolute LFC greater than 0.25, of which 218 were downregulated and 259 were upregulated following (table S3). Among the upregulated genes, we identified several well-known pro-inflammatory genes, such as *IL6* (LFC = 27.06), *IL1A* (LFC = 4.02), and *CCL4* (LFC = 3.27) (Fig. 4D). NE treatment resulted in 319 DEGs, with 209 downregulated and 110 upregulated (table S4). Similar to CORT exposure, we observed the upregulation of well-established inflammatory cytokines and chemokines, including *CXCL1* (LFC = 2.46), *IL1B* (LFC = 3.21), and *CXCL8* (LFC = 2.41) (Fig. 4E). These genes—particularly *IL6*, *IL1B*, and the chemokines *CXCL1* and *CXCL8*—are central mediators of inflammation, and their dysregulation has been implicated in the pathogenesis of various inflammatory and autoimmune diseases, highlighting the potential impact of stress-induced immune modulation.

To validate the enrichment of inflammatory gene expression in *CD163*⁺ cells following cortisol exposure, we performed RNAScope analysis looking specifically for cells expressing *CD163* that also express *IL6* and *IL1A*. We observed an increase in the number of *CD163*⁺ cells expressing *IL6*, as well as a marked increase in *IL6* expression within *IL6⁺ CD163*⁺ cells, as evidenced by a higher number of *IL6* puncta (Fig. 4F & G). Additionally, while the proportion of *CD163*⁺ cells expressing *IL1A* did not change, there was a trend (p = 0.08) toward increased *IL1A* expression within these cells, as indicated by an increase in IL1A puncta (Fig. 4F & G). Taken together, these analyses identify *CD163+* MonoMacs as a novel stress-activated innate immune cell subtype and further characterize distinct stress-responsive, disease-relevant gene signatures within this cell subtype.

### *CD163+* MonoMacs-specific stress-inflammation transcriptional risk score is associated with disease

To explore the potential clinical and translational relevance of our findings, we sought to develop a transcriptional risk score (TRS) informed by our previous findings. By integrating the cumulative effects of multiple genomic sites, these composite markers are thought to better capture risk for complex diseases as compared to single risk genes (*60, 61*).

Building on this framework, we developed a Cell-type-specific (*CD163*+) Stress-Inflammation Score (hereafter denoted CytSIS). This parsimonious score was generated by identifying the CORT and NE DEGs in *CD163*+ cells that overlapped with the hallmark inflammatory response gene set (n = 10 genes), weighting each gene by its treatment effect size (summed regression coefficient across CORT and NE treatments), and summing the resultant weighted gene expression products (Fig. 5A-B; table S5). Application of CytSIS to our scRNA-seq dataset demonstrated that stress hormone exposure robustly increased CytSIS in *CD163*+ cells (F-value = 248.25 p-value = 1.18e-104) (Fig. 5C), as expected.

**Fig. 5.**
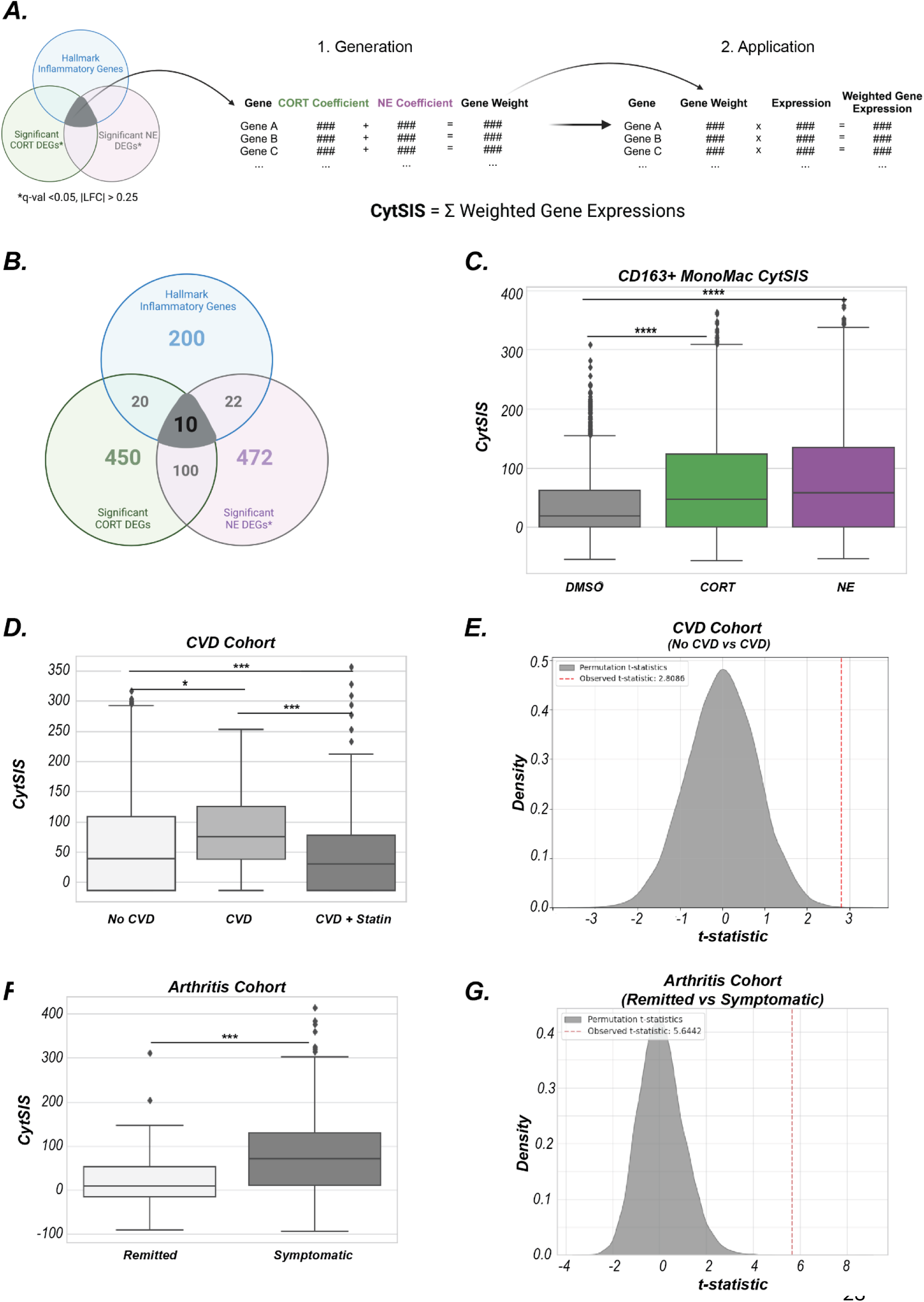
Cell-type-specific (*CD163+*) Stress-Inflammation Score (CytSIS) and its application in stress-associated diseases. (A) Overview of the methodology for generating the transcriptional risk score (TRS) CytSIS. The score was developed using *CD163*+ MonoMac inflammatory gene expression, and this score was applied across different single-cell RNA sequencing datasets, including those investigating stress-associated diseases. (B) Identification of 29 overlapping genes present in both the CORT and NE DEGs and the hallmark inflammatory gene set. These genes were used to generate CytSIS (Supp Table 5). (C) Application of CytSIS to our dataset shows significantly higher CytSIS scores in stress hormone-treated groups compared to control CD163+ cells, as expected. Statistical significance was determined by a One-way ANOVA and Tukey Post-hoc test. ****p < 0.0001. (D) Application of CytSIS to a cohort of HIV-infected individuals, categorized by cardiovascular disease (CVD) status. CytSIS scores are significantly higher in individuals with CVD compared to those without CVD, with scores decreasing in individuals with CVD who were treated with statin. Statistical significance was determined by a One-way ANOVA and Tukey Post-hoc test. *p < 0.05, **p < 0.01, ***p < 0.001, ****p < 0.0001. (E) 10,000 permutations were conducted, generating TRS by randomly drawing the same number of genes (n = 3) from the hallmark inflammatory genes and weighing them by the same treatment effect sizes. For each permutation, the TRS was then compared between the *CD163+* cells in the CVD and non-CVD groups. CytSIS significantly outperformed the random permutations. (F) Application of CytSIS to a cohort of individuals with rheumatoid arthritis, grouped based on symptom severity (remission, low, and moderate arthritis). CytSIS was significantly elevated in the low and moderate groups compared to the remission group. Statistical significance was determined by a One-way ANOVA and Tukey Post-hoc test. *p < 0.05, **p < 0.01, ***p < 0.001, ****p < 0.0001. (G) 10,000 permutations were conducted, generating TRS by randomly drawing the same number of genes (n = 10) from the hallmark inflammatory genes and weighing them by the same treatment effect sizes. For each permutation, the TRS was then compared between the remission and low arthritis groups. CytSIS significantly outperformed the random permutations.

To assess the applicability of CytSIS to cardiovascular disease, we next applied it in a cohort of PBMCs from HIV-infected individuals, classified into three groups: those with no cardiovascular disease (CVD), those with untreated CVD, and those with CVD treated with statins (*62*). The analysis focused on *CD163*+ cells (Fig. S5A). CytSIS was increased in the *CD163+* cells of the CVD as compared to the non-CVD and statin-treated CVD groups (F-value = 17.5440, p-value = 3.2606e-08; Fig. 5D). To confirm that this difference is driven specifically by the transcriptional effects of stress and is not due to non-specific differences in inflammation-related gene expression across groups, we conducted 10,000 permutations whereby we generated TRS by randomly drawing genes from the hallmark inflammatory gene list expressed in this cohort (n = 4), and weighing them by the scRNA-seq-trained effect sizes. For each permutation, the TRS was then compared between the CVD and non-CVD groups. CytSIS significantly outperformed the random permutations (p perm = 0.0003) (Fig. 5E). We observed similar findings when drawing from genes in the whole transcriptome (Fig. S5B).

We also applied CytSIS to a cohort of individuals with rheumatoid arthritis, who were grouped based on their symptoms into remitted or symptomatic arthritis (*63*). In *CD163+* expressing cells (fig. S5C), CytSIS was significantly elevated in the symptomatic compared to remitted group (T-statistic = 5.310, p-value = 1.356e-06) (Fig. 5F). We again performed 10,000 permutations of randomly generated weighted inflammatory TRS (n genes = 10) that were compared between remitted vs. symptomatic groups. CytSIS significantly outperformed the random permutations (pperm = 0.00) (Fig. 5G). We observed similar findings when drawing from genes in the whole transcriptome (Fig. S5D).

Together, these findings indicate that cell-type-specific measures of stress-related inflammation show promise as novel markers of disease risk.

## DISCUSSION

Prior work underscores immune dysregulation as a potential key process linking stress with disease, but how stress hormones affect distinct immune cells and cell types has remained unknown. This study leveraged single-cell transcriptomic signatures to determine how the stress hormones CORT and NE influence immune function and the relevance of these transcriptomic signatures to stress-related conditions.

Prior research has highlighted a complex relationship between stress exposure, stress hormones, and immune dysregulation in vivo (*64–67*). Both human and rodent studies have shown that stress exposure induces widespread shifts in immune cell types (*66, 68–70*). However, in vivo studies cannot disentangle whether these shifts reflect mere mobilization of particular cell types or reprogramming towards different immune cell types. Using our ex vivo that is devoid of the confounding effects of immune cell mobilization, we found that CORT exposure reduced plasmacytoid dendritic cells and CD8 naïve T cells (Fig. 2B). The decrease in CD8 T cells aligns with in vivo findings, where corticosterone administration in mice significantly reduced circulating CD8 T cells, suggesting cell death or identity changes (*71*). While corticosterone also changes the proportions of monocytes, neutrophils, and B cells in vivo (*71*), the lack of significant changes in these populations in our model indicates these shifts are likely due to cell mobilization only. Similarly, NE exposure decreased plasmacytoid dendritic cells and memory B cells while increasing CD4 T cells (nominally significant) (Fig. 2C). This aligns with in vivo findings showing a trend toward increased CD4 T cells after norepinephrine administration. This same in vivo study also demonstrated a dramatic increase in neutrophils following norepinephrine exposure, attributable to cell mobilization (*71*).

Our compositional analysis also revealed that the monocyte-to-lymphocyte ratio (MLR) increases following cortisol exposure. Elevated MLR has been reported in individuals experiencing high stress compared to those with low stress exposure (*64*), suggesting that cortisol may be a key driver of this pathology. Additionally, MLR has been studied as a marker of disease severity (*51–54, 72–74*), supporting a plausible link between stress and disease.

In this study, we demonstrate that stress hormones induce cell-type-specific transcriptomic changes, shedding new light that may help reconcile prior research supporting a range of pro- and anti-inflammatory effects of stress hormone exposure. For instance, some studies report that glucocorticoids reduce pro-inflammatory cytokines such as TNF-α and IL-1β, while increasing anti-inflammatory cytokines like IL-10 (*75*). Conversely, other studies suggest that glucocorticoids can promote the secretion of pro-inflammatory cytokines like IL-1β and IL-18 (*22*). Similarly, conflicting evidence exists regarding the inflammatory effects of NE (*76, 77*). Notably, our findings show that stress hormones can have opposite effects even within closely related cell subtypes (e.g., *VCAN*+ MonoMacs vs. *CD163*+ MonoMacs), highlighting the importance of studying distinct immune cell subpopulations in stress research. Nevertheless, we also identify a single gene differentially expressed consistently across all cell types for CORT (*FKBP5*) and another for NE (*RNU4-2*), suggesting that these genes could serve as immune markers of stress exposure.

Our inflammation analysis revealed that the effects of stress hormones are cell-type-specific, with the potential to either increase or decrease inflammatory gene expression depending on the cellular context. Notably, *CD163*+ MonoMacs exhibited a particularly significant increase in inflammation-related gene expression following exposure to both CORT and NE, highlighting their broad importance in stress-driven inflammatory states. This finding is intriguing given that *CD163*+ macrophages are widely described in the literature as having anti-inflammatory phenotypes, often referred to as alternatively activated macrophages (*78, 79*). However, it is important to note that *CD163*+ macrophages are also frequently observed in inflammatory and tumorigenic microenvironments, where they can contribute to disease-related inflammation through mechanisms distinct from those of classically activated macrophages (*80–83*). These cells are further studied for their pro-inflammatory and pro-atherogenic roles in atherosclerosis, where they may promote plaque progression and instability (*81–83*). Thus, our finding that CORT and NE induce inflammation-related gene expression in *CD163+* MonoMacs could provide a mechanistic link between environmental stress and highly prevalent conditions such as cardiovascular disease. This hypothesis is further supported by applying our transcriptional risk score, CytSIS, in cohorts of stress-associated diseases, highlighting the potential role of stress-induced inflammation in disease pathogenesis.

Our dataset provides a valuable resource for developing transcriptional risk scores (TRS), which are increasingly being used as biomarkers for disease risk and progression (*60, 61*). By leveraging cell-type-specific transcriptional responses to stress, we can create more precise tools for identifying individuals at risk of stress-associated diseases. For example, we demonstrated that *CD163*^+^ cell-specific transcriptional signatures effectively distinguish between healthy and diseased individuals. This approach provides a potential framework for leveraging single-cell datasets to promote early risk stratification and personalized intervention strategies. These findings may not only advance the development of disease biomarkers but also highlight potential therapeutic targets for mitigating the impact of stress and other environmental exposures on health.

## MATERIALS AND MEHTODS

### Ethics approval and participant consent

Human peripheral blood samples were obtained from healthy donors, aged 19 −28, at the UNC Blood Donation Center with informed consent and IRB approval. Eight healthy volunteers participated in the scRNA-sequencing study and q-PCR experiments, and four independent donors in the RNAscope experiments.

### Isolation of PBMCs and Neutrophils

PBMCs and polymorphonuclear neutrophils (PMNs) were isolated from human peripheral blood samples as previously described (*84*).

12 mL of fresh peripheral blood was collected using vacutainer EDTA tubes, diluted with 12 mL of DPBS supplemented with EDTA (2mM) and FBS (1%), and then gently layered over Lymphoprep (STEMCELL Technologies, USA) in a 50 mL conical tube. Samples were centrifuged at 800g for 15 min at room temperature with breaks off. After centrifugation, PBMCs in the white foamy layer were carefully transferred to a new tube. After being washed with DPBS twice, samples were then centrifuged at 250g for 10 min, and the pellets were resuspended with cell culture media (RPMI supplemented with 5% of human serum). Cell counts were performed on the automated cell counter and 250.000 cells/well were seeded into 24-well plates, incubated at 37C in a 5% CO2 incubator for 1 hour, and then treated with 100 nM Cortisol, 10 nM Norepinephrine, or 0.0001% DMSO (vehicle control) overnight.

Given their shorter lifespan in cell culture (*85*), PMNs were recovered from the red blood cells layer following 1 x RBS lysis buffer digestion procedure and washed twice with 1X PBS and resuspended in cell culture media. Cells were then seeded into 24-well plates at a seeding density of 250,000 cells/well, incubated at 37C in a 5% CO2 incubator for 1 hour, and then treated with 100 nM Cortisol, 10 nM Norepinephrine, or 0.0001% DMSO for 4 hours.

### Library preparation and sequencing

Cells were removed from the incubator, and the plates were centrifuged at 500g for 5 minutes for pelleting. Cells were washed with 1X PBS and carefully detached from each well by gentle pipetting, then resuspended in 1X PBS to create a single cell suspension. Cells were fixed immediately using a Fixation Kit (Parse Biosciences), counted, and stored at −80 °C as per the manufacturer’s protocol.

The single-cell libraries were constructed according to the Parse WT kit manufacturer’s protocol. 4166 cells per sample (for a total of 100,000 cells from 24 samples) were barcoded using the reverse transcription barcoding approach. Barcoded cDNA was amplified and verified via TapeStation for size distribution following fragmentation, end repair, A-tailing, and adaptor ligation to complete library construction. All libraries were sequenced as paired-end 150-bp reads using the Illumina NovaSeq X Plus 300 platform with a reading depth of 83-104M reads per sample.

### Preprocessing

Preprocessing was performed using Scanpy (v1.9.3) following a previously established workflow (*86*). Cells expressing fewer than 300 genes were excluded, as were genes expressed in fewer than 5 cells. To further refine the dataset, cells expressing more than 8,000 genes or with total gene counts exceeding 20,000—likely representing doublets—were removed. Additionally, cells with mitochondrial gene expression exceeding 15% of the total counts were filtered out to exclude unhealthy or dying cells. Clusters were annotated using the top DEGs, canonical markers, and reference data from previously published single-cell RNA sequencing datasets (*49, 50*). No size cutoff was used to remove small clusters. Highly expressed genes within each subset were identified using Wilcoxon rank testing implemented in Scanpy.

### Compositional Analysis

Cell densities in each subset were calculated and plotted for treatment samples versus controls using Scanpy embedding density functions (*86*). For each sample, the cell type composition was calculated as the proportion of cells from a specific cell type relative to the total cells in that sample. To minimize noise due to inter-individual differences and thus maximize statistical power, differences in cell type composition were assessed using paired t-tests, pairing each donor’s treatment sample with their corresponding control sample. Monocyte to Lymphocyte Ratio (MLR) was calculated by adding all cell types that could be classified as monocytes (CD163+ MonoMac, CD74+ MonoMac, IL7R+ MonoMac, VCAN+ MonoMac, CD14-MonoMac) and divided by the sum of all cells from cell types classified as lymphocytes (Nai CD4 T, Mem CD4 T, Nai CD8 T, Mem CD8 T, Th17, Nai B, Mem B, Plasma B). Statistical differences between control and treatment groups were calculated with paired t-tests. Two-sided *p* < 0.05 was the threshold for statistical significance.

### RNAScope

PBMCs were isolated from four healthy donors (two male, two female) and treated as previously described. Following overnight incubation, cells were collected into centrifuge tubes and centrifuged at 300 × g for 5 minutes. The resulting cell pellets were resuspended in 10% neutral buffered formalin (NBF) and fixed for 15 minutes at room temperature. After fixation, cells were centrifuged again at 300 × g for 5 minutes. The supernatant was carefully removed, and the pellets were resuspended in 20–30 μL of pre-warmed HistoGel (Thermo Scientific, HG-4000-012). While still in liquid form, the cell-HistoGel mixture was pipetted onto a parafilm to form a small gel droplet encapsulating the PBMCs. The droplet was allowed to solidify and then immersed in 10% NBF for an additional hour to ensure thorough fixation. Subsequently, the gel droplet was transferred to 70% ethanol and stored until further processing.

The gel droplet containing PBMCs was processed as formalin-fixed paraffin-embedded (FFPE) tissue and sectioned onto slides for RNAscope staining, following the protocol provided by Advanced Cell Diagnostics (ACD). Depending on the experimental conditions, cells were stained for either CD163, IL1A, and IL6, or CD3E, NELL2, and IL8A, along with DAPI nuclear counterstaining. Imaging was performed using a Zeiss LSM 980 confocal microscope equipped with a 64× oil immersion objective. Quantification of RNA signals was conducted using the ImageJ Cell Counter plugin, enabling the enumeration of individual transcript puncta per cell.

### Differential Gene Expression Analysis

Differentially expressed genes (DEGs) were identified using the rank_genes_groups function from Scanpy (*86*). The analysis compared the CORT and NE treatment groups to the control group using the Wilcoxon rank-sum test. Gene expression data from the raw count matrix was utilized for this analysis to ensure accurate assessment of gene expression differences. False discovery rate (FDR) correction was applied using the Benjamini-Hochberg method to account for multiple hypothesis testing. Adjusted p-values and log-fold change were used to rank significant DEGs.

To identify cell-type-specific gene expression changes, subsets of data from each cluster (cell type) were created, and the rank_genes_groups function was applied again to detect DEGs. The top 10 DEGs from the bulk analysis were then visualized based on log fold change to assess heterogeneity in differential expression across cell types (Fig. 3B and D).

### qPCR

Total RNA was extracted with an RNeasy kit (Qiagen, cat. No 74106) from cells exposed to DMSO, cortisol, and norepinephrine. The Super Script IV VILO cDNA kit (Invitrogen, cat.no 11756500) was used to synthesize cDNA. mRNA levels of the following geneswere assessed by q-PCR: VSIG4, FKBP5, ZBTB16, RNU4-2, CXCL1, IL1B.

Detailed information on Q-PRC primer sequences is provided later. Reactions were performed with two replicates in a 96-well plate using the Quant Studio 6 (Applied Biosystems). The housekeeping gene *GAPDH* was used as a reference control.

### AUCell

We employed the AUCell algorithm to score the activity of our gene subset of interest across individual cells, identifying those with significantly higher transcriptomic activity (*58*). This approach involves ranking all genes within each cell based on their expression levels, effectively creating a prioritized list of genes per cell. AUCell then calculates the Area Under the Curve (AUC) for each gene set within these rankings, providing a score that reflects the enrichment of the gene set among the top-expressed genes in each cell. This method allows for the identification of cells where specific gene sets are actively expressed, independent of the gene expression units and normalization procedures used. Given our focus on inflammatory gene expression, we selected the Hallmark Inflammatory Response gene set from Gene Set Enrichment Analysis (GSEA), consisting of 200 genes, as our gene set of interest (*59*).

After assigning each cell a single-gene enrichment score, we calculated the difference (delta) between the control and treatment groups within each cell type’s AUCell score. Statistical differences between control and treatment groups were assessed using Student’s T-test, with multiple testing corrected using the Bonferroni method.

### CytSIS

#### Development of the Cell-type-specific Stress-Inflammation Score (CytSIS)

Informed by our findings, we developed a transcriptional risk score (TRS) termed CytSIS. By integrating the cumulative effects of multiple genomic sites, this composite marker approach is thought to better capture risk for complex diseases as compared to single risk genes (*60, 61*). CytSIS was generated by identifying the differentially expressed genes (DEGs) resulting from CORT and NE treatments in *CD163*+ MonoMacs that overlapped with the Hallmark Inflammatory Response gene set (n = 10 genes) (Table S5). For each DEG, the treatment effect size was calculated by summing the regression coefficients from CORT and NE treatments compared to the control, as determined through the Wilcoxon rank-sum test. Each gene was thus weighted by its treatment effect size.

#### Application of CytSIS

To apply the CytSIS score, we first created a subset with cells in the cohorts that expressed the *CD163* gene. We then calculated the product of each individual gene’s weight (trained using our scRNA-seq treatment dataset) and its gene expression value for each cell (derived from the human cohort dataset). This calculation was performed only for the genes that were available in each dataset (4 genes for the CVD cohort and 10 genes for the arthritis cohort). The final CytSIS score for each cell was obtained by summing these products across all genes in the set, yielding a single score that reflects the relative stress-related inflammatory activity of the cell.

#### Permutation Testing

To validate the specificity of the CytSIS score, permutation testing was performed to rule out the possibility of the score being driven by random gene expression patterns. For each cohort (CVD and arthritis), 10,000 permutations were conducted where a random selection of genes (matched to the number of genes available in each dataset: 4 for CVD and 10 for arthritis) was drawn from the hallmark inflammatory response gene set (or the whole genome) and weighted by the same treatment effect sizes. The resulting permuted scores were then compared to the CytSIS scores to assess whether CytSIS outperformed the random permutations. Statistical significance was determined by comparing the observed CytSIS score to the distribution of the permuted scores.

### Cohorts

Human cohorts used for CytSIS testing were obtained from publicly available data repositories.

#### Cardiovascular Disease (CVD) (GEO: GSE205320)

All participants in this study were part of the Women’s Interagency HIV Study (WIHS) (*62*). From the initial 1,865 participants in the WIHS vascular sub-study, 32 participants were selected for scRNA-seq analysis of PBMCs. The presence of carotid artery focal plaque was used to classify participants into four groups of eight participants: HIV−, HIV+CVD−, HIV+CVD+, and HIV+CVD+ on cholesterol-reducing treatment (CRT). For our analysis, we focused on the role of stress in CVD and, to rule out confounding by other disease contributions, we excluded the HIV− group.

#### Rheumatoid Arthritis (RA)

Participants from this study consist of patients meeting the American College of Rheumatology (ACR) classification criteria and were recruited from UCSF rheumatology between 2016 and 2020 (*63*). PBMCs were collected for scRNA-seqThe disease Activity in 28 joints using CRP (DAS28-CRP) was used to stratify the patients in remitted (n = 2), or symptomatic (n = 14) groups with low or moderate disease activity according to the 2019 updated ACR recommendation on disease activity measures (*87*).

## Acknowledgments

The authors acknowledge Laura Huff for her valuable feedback on this manuscript. We also thank Dr. Tessa-Jonne Ropp of the UNC Neuroscience Microscopy Core for her assistance with image acquisition, and Ashley Ezzell for her guidance in developing the methodology for PBMC RNAscope experiments.

## Author contributions

Conceptualization: SB, OK, AZ

Methodology: SB, OK, LC, NS, BM

Investigation: SB

Visualization: SB, LC

Funding acquisition: AZ

Project administration: SB, AZ

Supervision: AZ

Writing – original draft: SB, OK

Writing – review & editing: SB, AZ

