## Supplemental Figures for "Single-cell transcriptomics yield insights into the stress-immune interplay and inform disease risk"

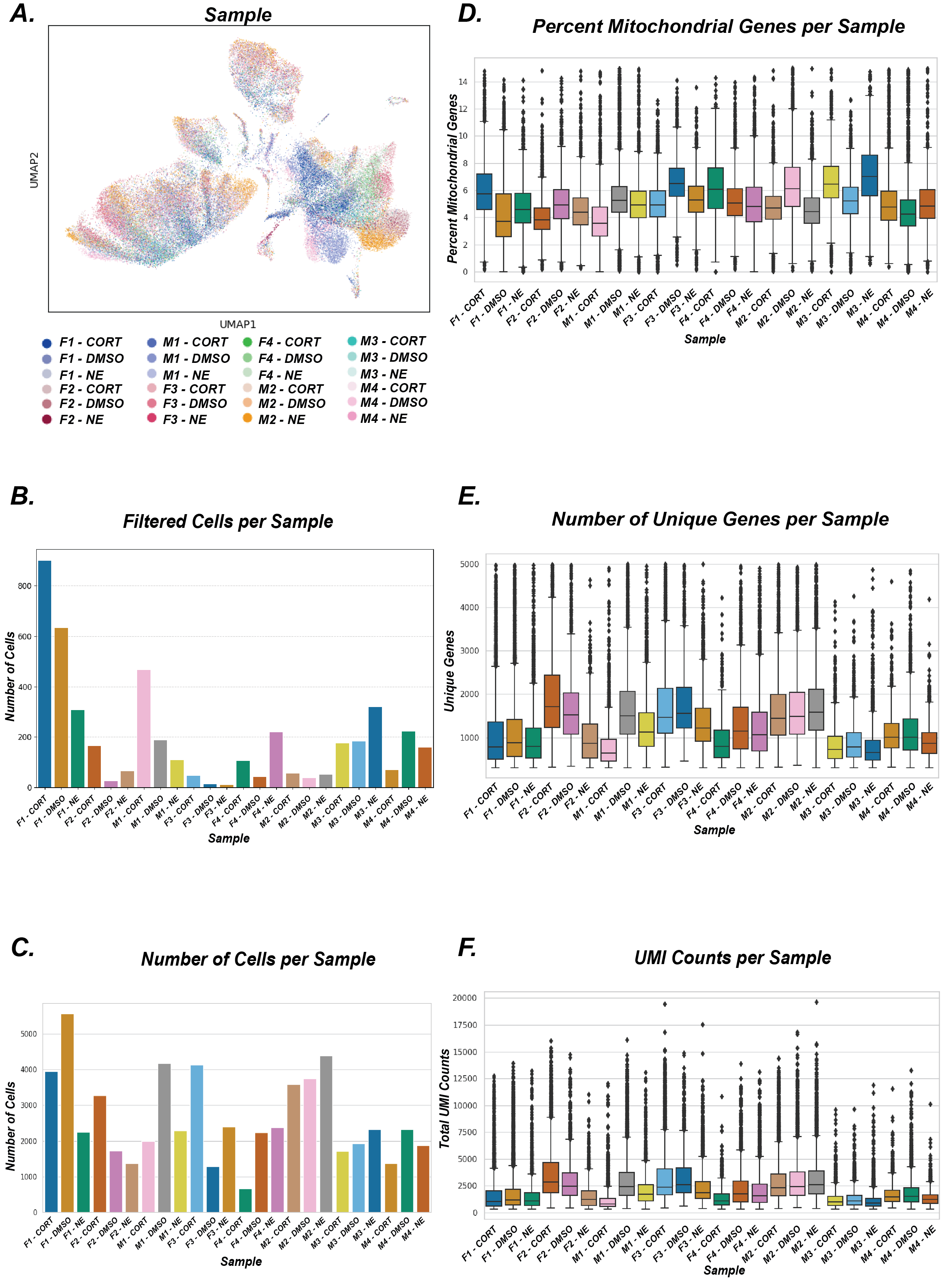

**Supplemental Fig 1. Overview of cell filtering and quality control metrics.**

(A) UMAP plots of all retained cells after filtering low-quality cells, showing distribution by sample.
(B–F) Sequencing and quality control metrics per sample after filtering.

(B) Number of cells filtered out per sample.

(C) Number of cells retained per sample.

(D) Percentage of mitochondrial gene expression per cell, per sample.

(E) Number of unique genes detected per cell, per sample.

(F) Total number of unique molecular identifiers (UMIs) per cell, per sample.

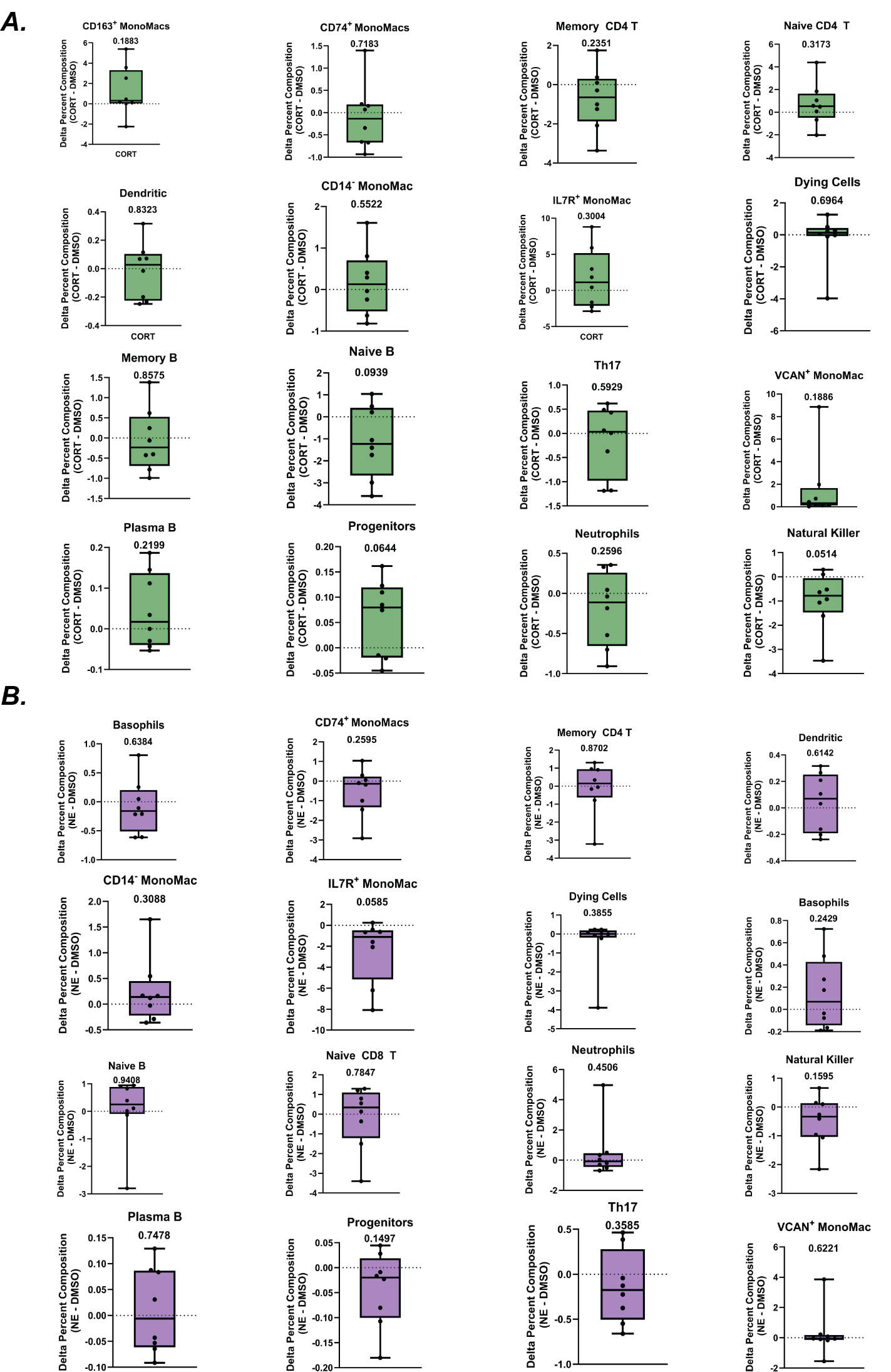

**Supplemental Fig. 2. Cell-type composition changes due to hormone treatment.**

(A) Changes in the percentages of all cell types following CORT treatment that did not reach statistical significance. The graphs show deltas between CORT and DMSO. Paired t-tests were used to test for significance.
(B) Changes in the percentages of all cell types following NE treatment that did not reach statistical significance. Graphs show the deltas between NE and DMSO. Paired t-tests were used to test for significance.

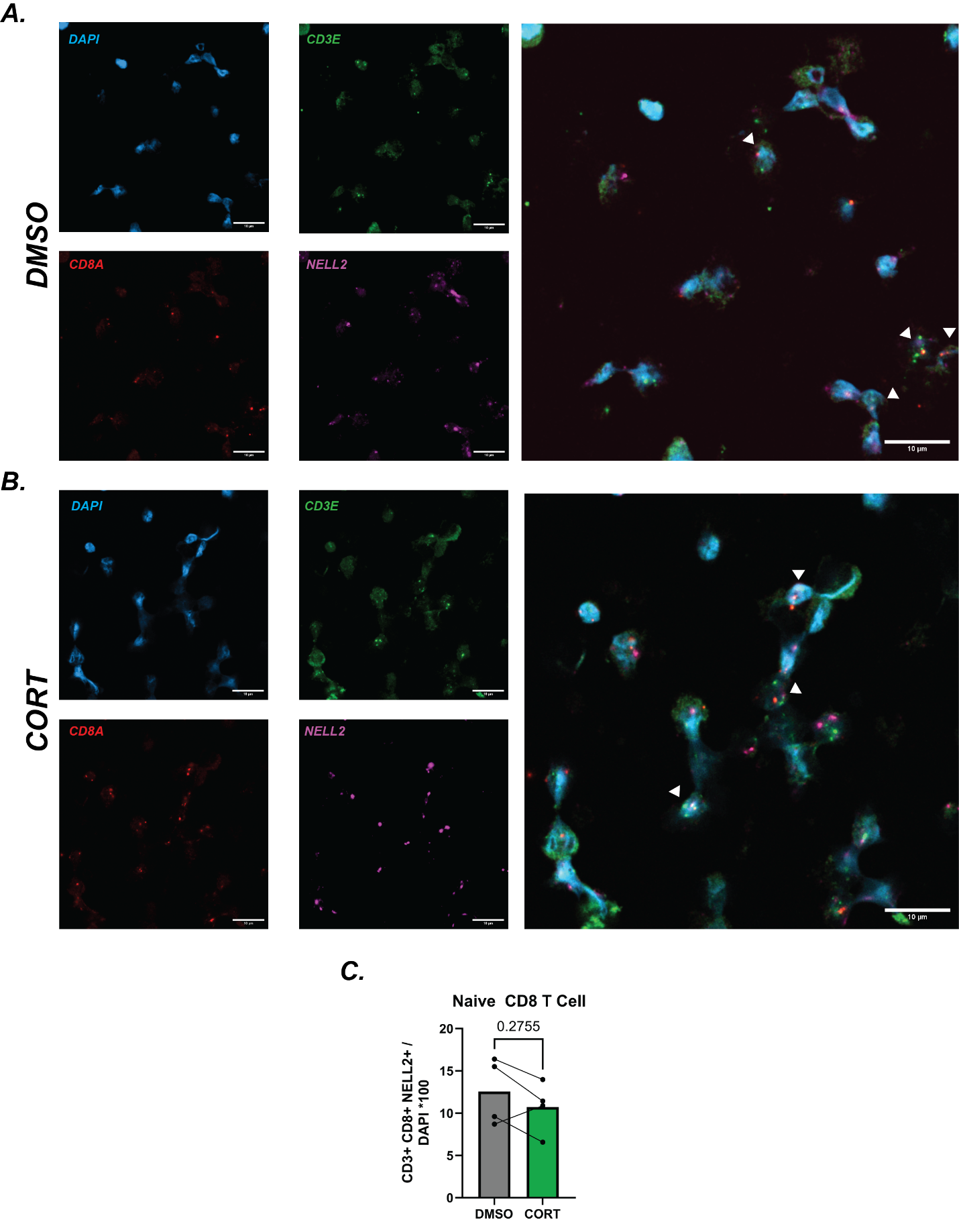

**Supplemental Fig. 3. Naïve CD8⁺ T cell composition changes following cortisol treatment.**

(A) Representative RNAScope image of control cells. Naïve CD8⁺ T cells are identified as triple-positive for CD3, CD8, and NELL2.
(B) Representative RNAScope image of cortisol-treated cells. Naïve CD8⁺ T cells are identified as triple-positive for CD3, CD8, and NELL2.
(C) Quantification of naïve CD8⁺ T cell proportions across donors. Statistical significance was assessed using Student’s *t*-test.

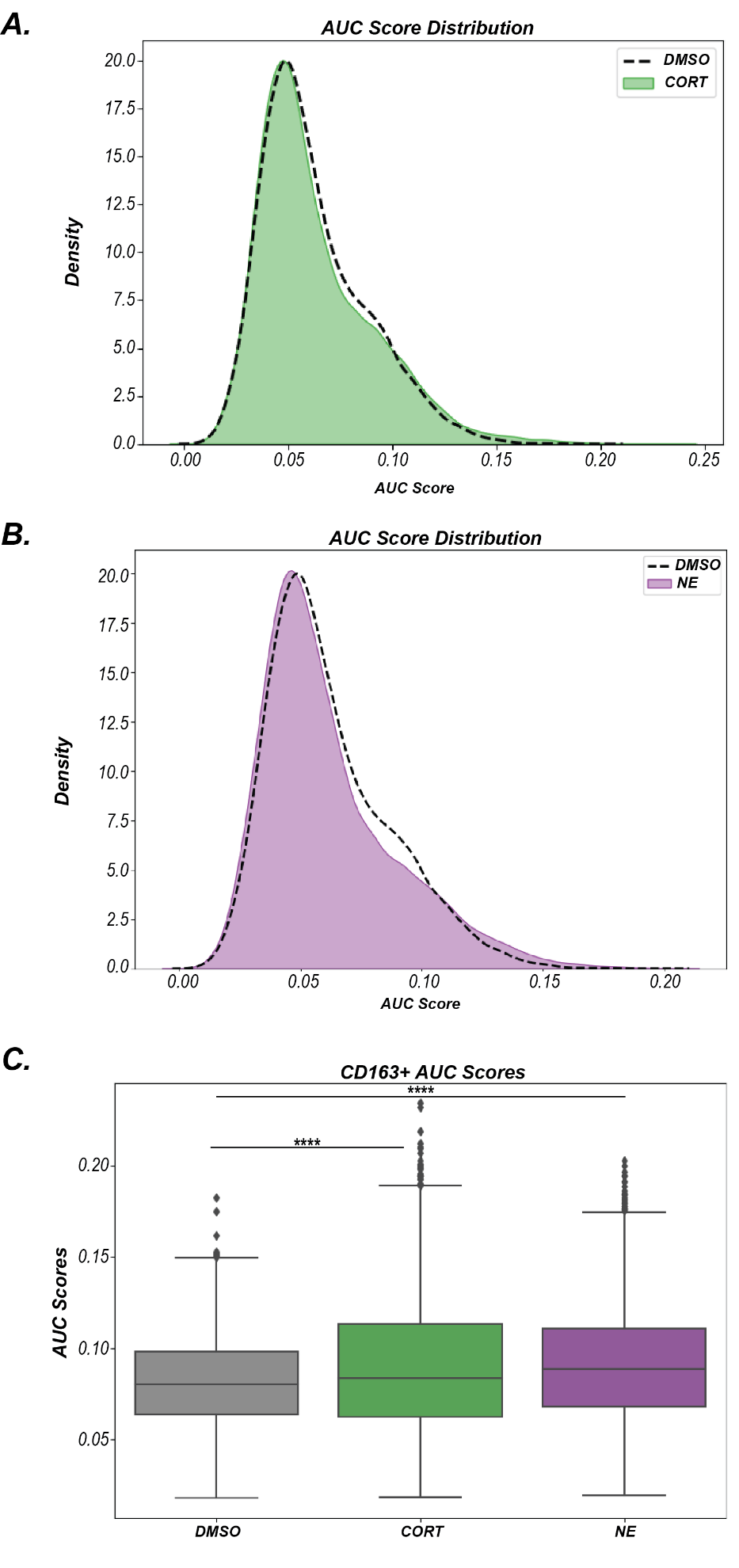

**Supplemental Fig. 4. Inflammatory gene expression changes due to treatment.**

(A) Distribution of cells based on their AUCell scores, comparing DMSO control (dashed line) and CORT-treated cells (green area).
(B) Distribution of cells based on their AUCell scores, comparing DMSO control (dashed line) and NE-treated cells (purple area).
(C) One-way Anova comparing AUCell score in *CD163+* MonoMac cells across treatment groups. Statistical significance was determined by a One-way ANOVA and Tukey Post-hoc pairwise comparison tests. ****p < 0.0001.

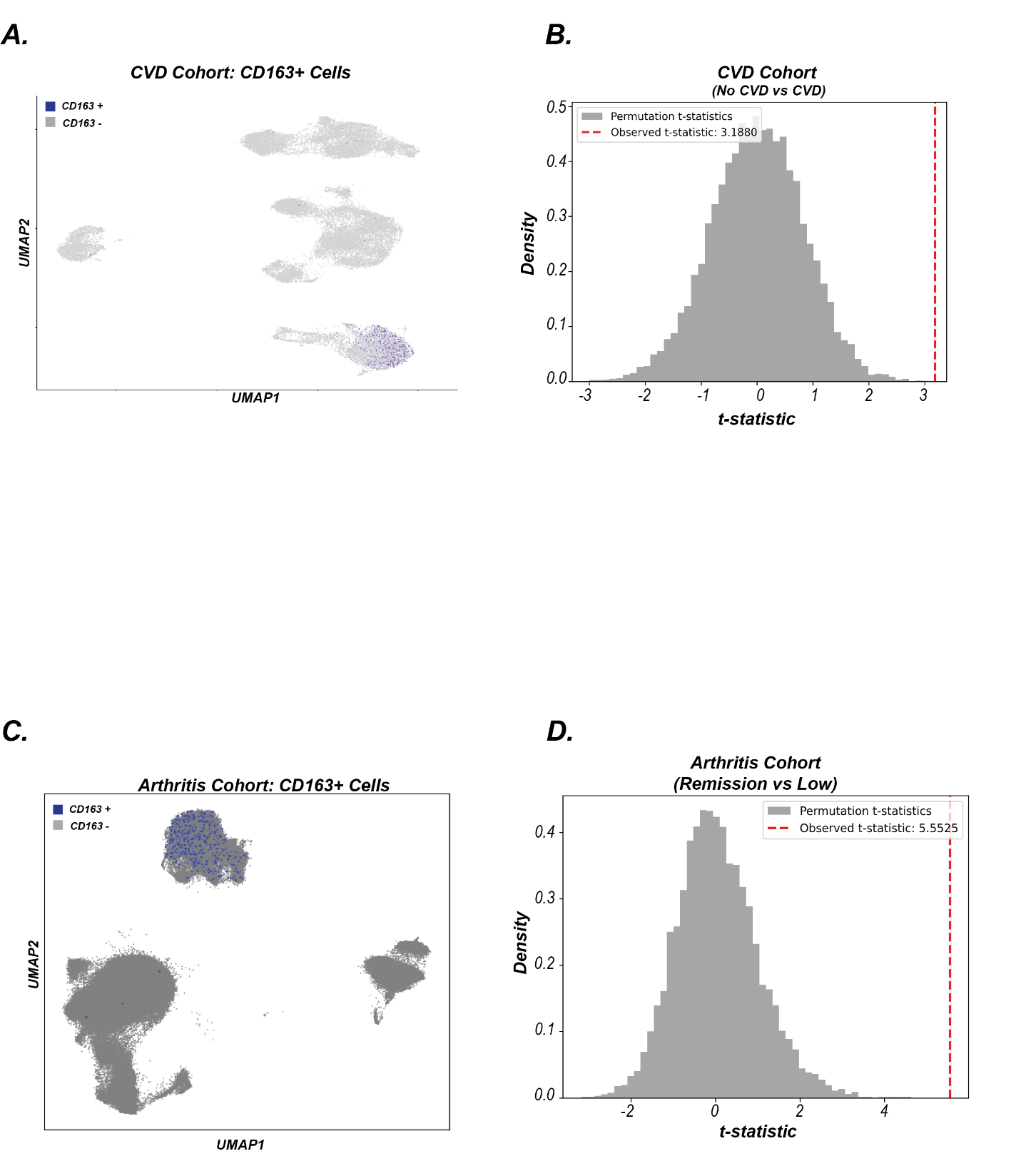

**Supplemental Fig. 5. CytSIS application in cohorts of stress-associated diseases.**

(A) UMAP visualization from the CVD cohort, highlighting CD163+ cells in blue.
(B) CVD cohort: 10,000 permutations were performed, each generating a transcriptional risk score (TRS) randomly selecting the same number of genes available in the dataset and used for CytSIS (n = 7) from the genome and weighting them by the scRNA-seq-trained treatment effect sizes (regression coefficients). The TRS was compared between the CVD and no CVD groups for each permutation. CytSIS significantly outperformed the random permutations (pperm = 0.00).
(C) UMAP visualization from the Arthritis cohort, highlighting *CD163+* cells in blue.
(D) Arthritis cohort: 10,000 permutations were performed, generating TRS by randomly selecting the same number of genes from the genome available in the dataset and used for CytSis (n = 29) and weighting them by identical treatment effect sizes. The TRS was compared between the remission and low arthritis groups for each permutation. CytSIS significantly outperformed the random permutations (pperm = 0.0003).
